# The Vacant Niche Revisited: Using Negative Results to Refine the Limits of Habitability

**DOI:** 10.1101/2023.11.06.565904

**Authors:** L.E. Ratliff, A.H. Fulford, C.I. Pozarycki, G. Wimp, F. Nichols, M.R. Osburn, H.V. Graham

## Abstract

To define the boundaries of habitability, biologists often search for highly specialized organisms in extreme environments. However, negative life detections—when a method is unable to detect microorganisms in a given setting—are just as important to constrain the environmental limits of life on Earth. In turn, these limits inform the selection of targets for life detection on other worlds.

We performed a comprehensive, though non-exhaustive, literature search for negative life detections in polyextreme environments. We then catalogued the physicochemical conditions at these sites to further understand the habitability limits for life on Earth and the effects of multiple stressors on habitability. Using multivariate statistical techniques, our study searched for combinations of environmental parameters where extremes support or inhibit life. Our search raised several methodological and analytical considerations relevant to life detection studies in extreme environments. Incomplete documentation of environmental factors and experimental protocol limitations in the extreme environment literature complicated our analyses. This demonstrates the need to report negative results, particularly in life detection experiments, and the potential value for standardized reporting protocols. Exploring the range of results possible from life-detection methodologies is key to constrain the limits of life on Earth and informs our search for life elsewhere.

## INTRODUCTION

The true limits of habitability can only be identified by studying locations where life is present *and* where it is absent. Habitability can be defined as the combination of physical and chemical conditions that sustain organisms^1, 2^. Typically, limits to habitability have been determined by using the presence of organisms in an environment as justification to push the limit “outward” toward an extreme, relative to average conditions experienced by humans and model organisms. As scientists have identified and cataloged microbial communities in increasingly extreme environments on Earth, the physical and chemical conditions under which life is known to inhabit have rapidly expanded^3^. This outward approach—searching for life by pushing towards the uninhabitable extreme conditions—can be supplemented by “inward” boundary-setting, which begins outside environments where life is found (Figure 1). Unlike literature detailing the discovery of microbes in increasingly extreme environments, an accounting of the number and persistence time of uninhabited environments has not emerged. Cataloging and organizing data on these environments helps constrain the limits of habitability and encourages more discussion of these understudied locations. These physicochemical limits to life on Earth can constrain and inform the search pattern for life on other worlds and facilitate exploration of the environmental changes that promote habitability^3, 4^.

**Figure 1.**
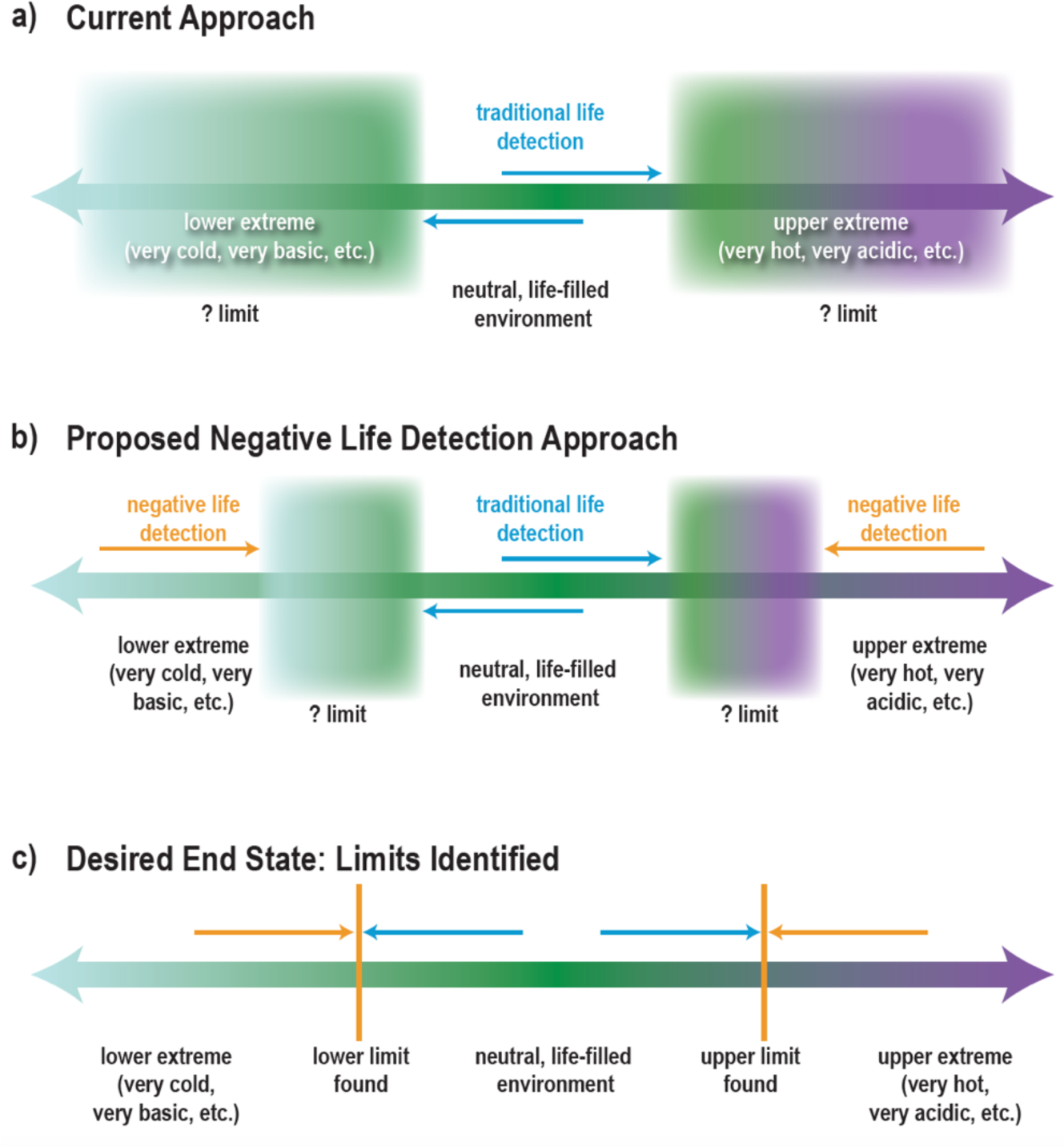
This study seeks to determine habitability limits by combining traditional and negative life detection approaches. Conventional life detection studies have typically explored the ranges of habitability by moving across a parameter gradient from moderate conditions into more harsh extremes, as exemplified by the blue arrows in (a) which point “outward” along a line representing the parameter space for a given variable. This strategy has identified life’s ability to contend with extremes but rarely are the conditions that truly limit life fully delineated. To complement the approach, negative life detection (b) seeks to determine the conditions under which life may not be found, allowing constraint of the unknown habitability limit for the given parameter. This is shown by the orange arrows pointing “inward,” creating an outer limit and then lowering that threshold. The threshold itself is identified by the gradient band, in which the limit resides but is unknown. With both of these methodologies together, the lower and upper limit can be constrained and potentially found, shown in (c).

Environments may be uninhabited because of physicochemical factors which inhibit survival, mechanisms that limit dispersal of organisms to those areas, or both. An uninhabited niche has a resource set and/or a physicochemical environment that is irreconcilable with the survival of life on Earth. On the other hand, a vacant niche is one which does contain potentially usable resources, *i.e., an uninhabited yet habitable environment*^5^. In fact, vacant niches do exist, and the conditions that are likely to form them overlap with the conditions which create extreme environments. A vacant niche may have either not been inaccessible due to dispersal barriers or organisms have yet to adapt mechanisms to contend with limiting physicochemical factors^6^.

There exists within environmental microbiology and astrobiology an optimistic tendency to assume that microbes are ubiquitous in every habitat on Earth, and that for every exotic environment or potential habitat there is an even more inventive metabolism that can take advantage of that niche^7^. With this perspective, failure to detect life is generally attributed to limitations on life detection methods rather than a significant finding. The fact that there can be multiple causes for an uninhabited niche complicates the use of negative life detection as determinant for habitability limits. As well as physicochemical parameters, quantification of factors that drive dispersal of individual microorganisms and communities, such as wind, water, and animal movement, can help assess whether dispersal is likely to have occurred and thus whether a niche is truly life-limiting or conditionally vacant^8^.

This study surveyed the relevant literature for results where a life detection experiment was performed but no life was detected at a given sampling location. By refining Terran habitability across multiple parameters, this study is aimed to assist efforts to develop an operational definition of life based on biochemical reactions that does not presuppose specific biochemistry or adaptation. In other words, understanding the true limits to life in extreme environmental conditions on Earth is necessary to postulate about the limits of life elsewhere in the universe.

## RESULTS

### Distribution of negative life detections

Our methodology identified 212 negative life detections out of 497 total samples catalogued. Sample sites can be categorized into 13 globally distributed biomes (Figure 2). Analyses included samples from high altitudes and the deep subsurface, desiccated terrains and aqueous solutions, pH values that span the 14-point scale, and temperatures ranging from -25 to 125°C. To quantitatively represent the diverse features of these environments, the database included 230 distinct physical, elemental, and chemical parameters overall (Table 1).

**Figure 2.**
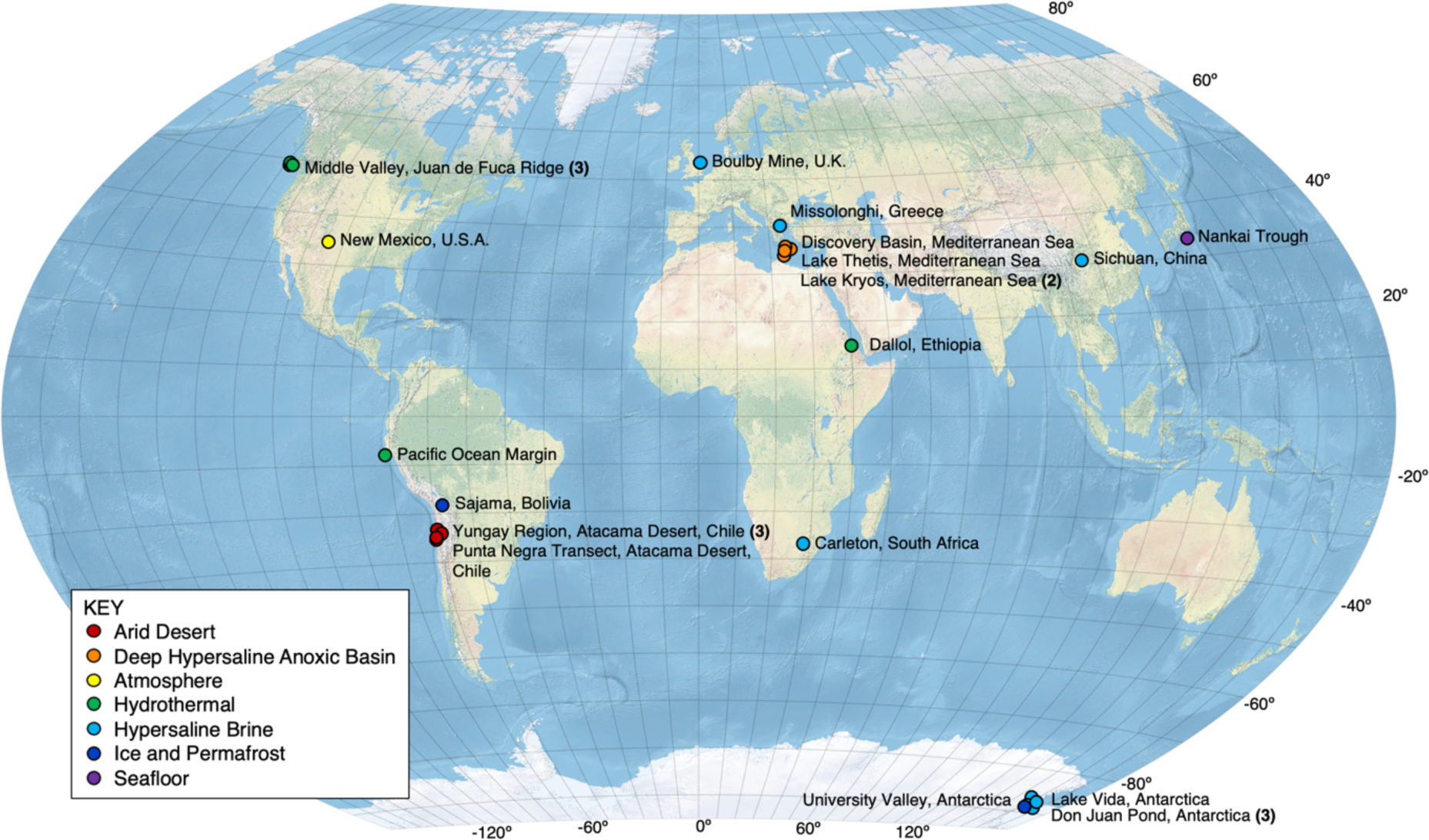
Negative life detections were cataloged from a wide range of potential biomes with vastly different physical and chemical conditions, occurring in aquatic and terrestrial settings. These sampling sites are often closely co-located with fully inhabited niches. The color of the dot identifies its biome genre. Where there are multiple dots overlapping, the number in parentheses after the location name indicates the number of separate sites.

**Table 1.** A demonstrative subset of the physicochemical parameters included in the database for this study, drawn from data available in the selected papers.

| Physical | Elemental | Chemical |
| --- | --- | --- |
| coordinates, pH, temperature, pressure, water activity, distance above/below sea level, density, season, buffer capacity, osmotic pressure, freezing point, solubility | Al, As, Au, B, Ba, Br, Ca, Ce, Cl, Co, Cr, Cs, Cu, Fe, Ga, Hg, K, Kr, La, Li, Mg, Mn, Mo, Na, Ni, Rb, P, Pb, S, Sb, Sc, Se, Si<sup>+</sup>, Sn, Sr, Th, Ti, Tl, U, V, Xe, Z | TOC, TIC, TON, TIN, DO, Eh, EC, ORP, C<sub>2</sub><sup>+</sup>, CaCO<sub>3</sub>, CO<sub>2</sub>, CO<sub>3</sub><sup>2-</sup>, CH<sub>4</sub>, HCO<sub>3</sub><sup>-</sup>, HS, H<sub>2</sub>S, H<sub>3</sub>BO<sub>3</sub>, KCl, NH<sub>4</sub>, NO<sub>3</sub><sup>-</sup>, PO<sub>4</sub><sup>-</sup>, SiO<sub>2</sub>, SO<sub>4</sub><sup>2-</sup>, stable isotope analysis, chaotropicity, kosmotropicity |

### Parameter space explored

Our survey recorded and collated all of the physical and chemical measurements made at each sampling site where a negative life detection was recorded. This includes parameters at both landscape- and sample-scale for chemical measurements ranging from elemental abundances to the characterization of soluble and insoluble organic species, metabolites, and nutrients. In addition to these physical and chemical factors, this study recorded negative or positive life detection outputs for the 42 unique life detection methods used in the surveyed studies. These included life-detection techniques using physical, chemical, and biotechnological methods, many of which were employed in tandem. Examples of these life detection methods include stable isotope analysis, organic geochemical analyses, and metagenomic data as well as imaging, microscopy and attempted culturing experiments (Figure 3).

**Figure 3.**
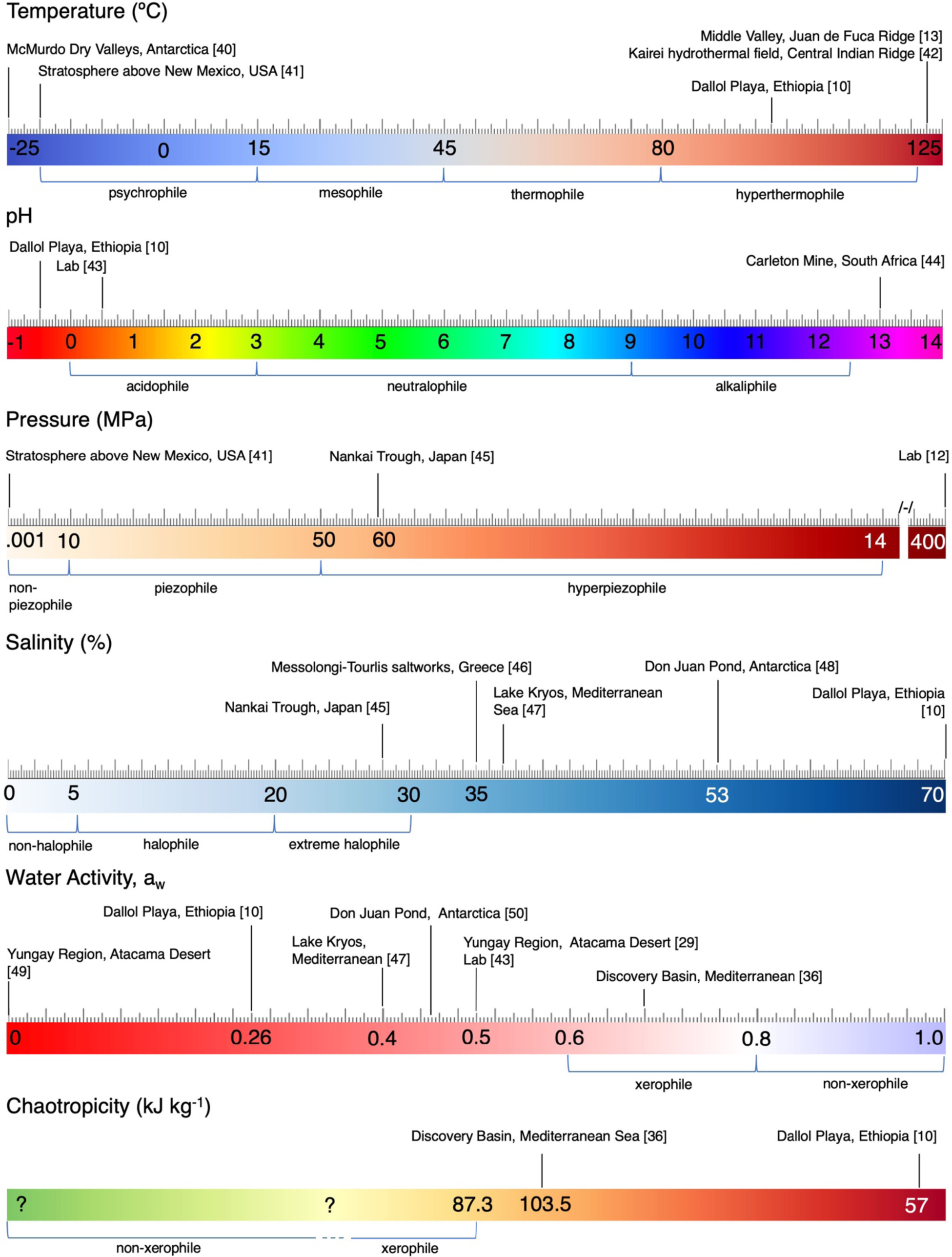
Various physicochemical extremes drive negative life detections. Six physicochemical parameters are shown with selected values for negative life detections from the literature. Citation numbers correspond to a negative life detection, reported as wholly or partially caused by that physicochemical factor and identified by the location of the sampling site. In the case of multiple negative life detections in one paper, the value depicted is the least biologically extreme. Brackets beneath each scale mark the widely accepted ranges for non-extremophiles and extremophiles of that physicochemical factor^3^. Scales are independent, without implied relationships between them. Chaotropicity is difficult to empirically calculate, making numerical determination of ranges challenging.

### Statistical analysis results

NMDS and ANOSIM analyses show that the observations in the data set are significantly differentiated by their biome genre, geographic location, and source study with a significance value of less than 0.05. The analyses revealed, however, that there was no statistical significance linking physicochemical parameters with life status (Figure 4, SI). The grouping by biome genre seen in the NMDS corroborates the ability of the NMDS to properly group the observations in ordination space, as environments within the same biome should demonstrate similar patterns between their physicochemical parameters. Within these clumped biome genres, both positive and negative life detections occur within close ordination space proximity across many different environments; there are even cases in which the points are superimposed, indicating the similarity of the sample environmental conditions for that sample.

**Figure 4.**
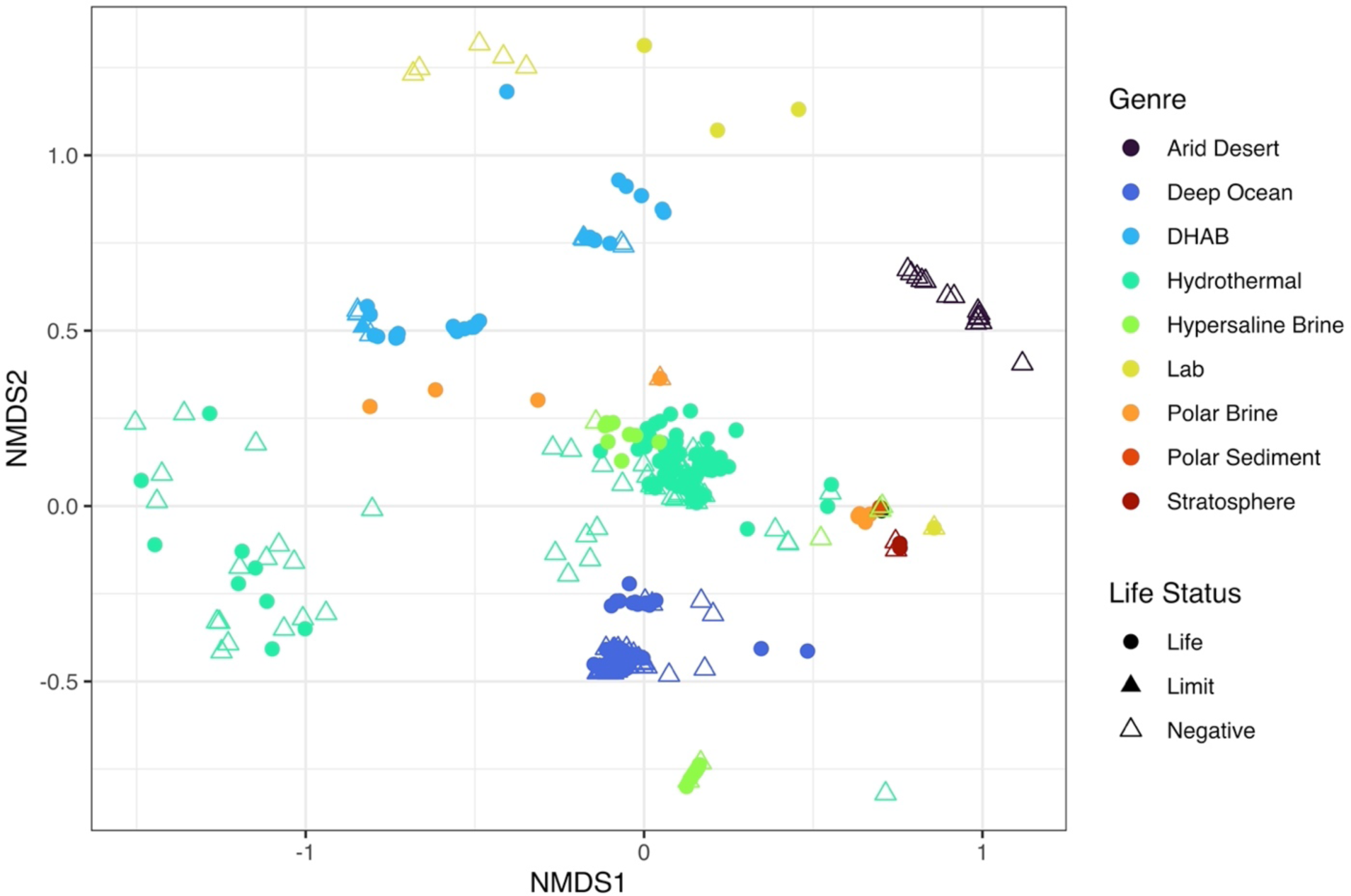
Physicochemical data plotted using NDMS shows clumping by biome genre, but does not group based on the outcome of the life detection effort. Each point represents all of the physicochemical parameters associated with one life detection sampling. The position of the points in two-dimensional space is representative of the similarity between each set of parameters describing a point’s environment. Points that are closer together have more similar physicochemical values, which explains the genres clumping together. Genres are denoted using color. The life status, denoted by the shape of the point, shows intermingling between life and negative points.

Bivariate logistic regression showed significant (p < 0.05) correlations between the probability of life in a sample and nine physicochemical variables (Temperature, pH, Water Activity, Distance Below Sea Level, Density, Alkalinity, and Barium, Magnesium, and Nickel concentrations), but none of the variables appear exclusively responsible for defining habitability within a location (see SI). For example, with increased water activity there was an increase in the presence of life. Below a water activity of 0.45 only those without life (negative results) are found. As with the NMDS ordination plot, there is a wide range of water activities (0.45 - 0.9) where both positive and negative detections are identified, indicating that water activity itself does not drive life status in this range.

Due to the paucity of studies that span the boundary of positive and negative life detection and the variety of different environments included in this study, the logistic regressions, while statistically significant, do not completely inform the limits of life. Our statistical analysis is complicated by negative life detections in polyextreme environments. Habitability is defined by more than one limiting parameter, even when one parameter would have constituted a formidable barrier. Isolating one physicochemical factor from a polyextreme environment to perform statistical analysis renders the logistic regression less meaningful.

### Polyextreme environments

A combination of extreme environmental parameters may together apply intolerable stresses to an organism or enable life outside the bounds of each constituent parameter. These may be referred to as co-limiting or co-enabling parameters, relying on co-adaptive mechanisms, depending on whether a set of multiple extremes experienced together restrict or facilitate life, respectively^9^. For example, low pH and high temperatures in surface water in the Dallol-Danakil area, Ethiopia, create seemingly vacant niches even though acidophiles and thermophiles grow at much lower pH or higher temperature when other physical factors are less extreme^10^.

Conversely, enzyme functionality increases at low temperatures and high pressures in high-salinity environments where exposure to one of these extreme parameters alone—either cold, pressure, or hypersalinity—causes denaturation^11^. These capabilities may arise from adaptations to one environmental factor which facilitate survival under other limiting factors^12^. This study intended to explore co-limiting and co-enabling factors with the negative life dataset. However, the data lacked the necessary coverage to conduct a statistically significant correlation between tolerance for multiple extremes. Future work could evaluate larger datasets of life status detections to identify physiological adaptations that confer tolerance in a polyextreme environment.

## DISCUSSION

Unique environmental challenges and experimental design limitations hinder the characterization of extremophile life and discovery of life’s limits. When microbial communities inhabit tenuous and/or fluctuating niches, environmental factors may prevent full identification of the microbial community studied. Additionally, choices made in protocol development—such as life detection technique choice and recorded variables—can shape the data’s interpretation and its broader comparability within the literature. This section will enumerate challenges that became apparent during the literature review and statistical analysis. The next section will recommend ways to address these challenges and improve the utility of data from extreme environments.

### Environmental Variables that Confound Life Detection Strategies

#### Low-biomass detections

Two factors complicated the assignment of life status to sampling from low-biomass areas. Firstly, the environments in this meta-analysis tended to contain gradients that approached the limits of life rather than supporting drastically different population sizes within close proximity. Ultimately, this means that many potential negative life detection sites are situated near to low-biomass areas. Secondly, life detection methodologies all have some level of noise and inaccuracy in their data, meaning that they may under or overreport the presence of life. Together, these facts create uncertainty in life determinations in low-biomass areas. While some negative determinations had no detected microorganisms, the study included some that had nonzero organisms but were below the confidence value of detection for the methodology used. For example, Cragg and Parkes^13^ and Cragg et al.^14^ calculated a “significance level” for their bacterial cell counts which was used as an upper cutoff for negative life detections from those studies. Cutoff values between low biomass samples and negative life detections, however, are not consistent between methodologies. Some quantitative life detection methods are more able to distinguish a true “negative” from a “near negative” than others, which makes life detection results difficult to compare between studies.

#### Indigeneity of life detections

In extreme, low-biomass sites or vacant niches, life detection methods may struggle to distinguish between indigenous, thriving communities and cells or cellular material that is preserved. Natural processes like aeolian transfer or gravitational particle movement in a water column may introduce foreign microbes which immediately perish. However, their genetic or metabolic materials may be preserved by low temperatures, high ionic strength, anoxic conditions, and protection from enzymatic degradation^15^. RNA and ATP, while less stable than DNA, can also be preserved in extreme environments. Ancient RNA signatures have been recovered from marine diatoms and plants ranging from 30,000 to 2.7 million years old, suggesting that the molecule may be much more stable than previously considered^16^. Similarly, high concentrations of particulate ATP are preserved in the anoxic brines of the Gulf of Mexico even though the microbial community in these brines is reportedly inactive^17, 18^. While the detection of biomolecules such as RNA and ATP is widely considered to be evidence of active cells, their preservation in extreme environments invokes a need for caution^19, 20^.

During sampling activities, researchers may inadvertently introduce non-indigenous microbes into the environment^21^. For example, Lake Vostok, a reportedly “pristine,” subglacial Antarctic lake, has been the focus of debate after Russian drilling activities may have introduced microbes into the environment^22^. The records held in ice and snowpack in the Antarctic are increasingly affected by motorized exploration activities and the presence of crewed field stations^23, 24^. Likewise, reports of microbial growth in the incredibly challenging conditions at Don Juan Pond may be the result of human activity and windblown contaminants^25, 26^. Both forms of contamination may alter the microbial abundance and composition of a sampling site.

An organism may be indigenous to an ecological niche if its optimal growth conditions reasonably match the ambient conditions of that site. Indigeneity could be addressed using a metastable equilibrium model which examines whether the proteins produced by strains present under extreme conditions have adaptations to lower energetic costs^27^. Just as microbial indigeneity can be evaluated at the cellular level, this method examines whether the proteins produced by microbes under extreme conditions have adaptations to lower energetic costs. These analyses can be carried out via genetic sequencing and translation of the proteins present to determine their structural adaptation^28^.

#### Effects of spatial and temporal scale

The physical scale at which a life detection is made influences both its determination and its extrapolative power to the site as a whole and to other environments with analogous physicochemical conditions. The challenges of instrument resolution with low biomass samples apply similarly when the sampled area is small and can lead to false negatives. Small-scale sampling may miss communities due to patchiness (environmental heterogeneity). The habitability of extreme environments can vary significantly on a scale of millimeters; for example, microbial community reports included in this study vary greatly between small gypsum deposits in the Atacama Desert^29^. Both sampling small areas and increasing the scale of experiments—at the cost of resolution—can convey an inaccurate sense of homogeneity across an environment. This makes the choice of sampling scale crucial in life detection, particularly in oligotrophic or low biomass regions^30, 31, 32, 33^.

Seasonal effects in an environment cause inconsistencies and knowledge gaps as well. Data collection often occurs over relatively short periods of time that may not capture the full scope of environmental conditions and population dynamics, particularly for species with slow division times. The driest sections of the Atacama Desert, such as the Yungay region, typically only experience rainfall for ten hours in an entire year^29^, and Western Australian transient lakes have distinctly different geochemical properties in dry and wet seasons. These desiccated microbial habitats are drastically different when exposed to precipitation, but sampling schedules may not be able to respond to rainfall. Likewise, the McMurdo Dry Valleys cannot be studied as extensively during austral winter due to harsh conditions and research restrictions, limiting our understanding of habitability at these sites.

### Experimental Design and Interpretation Challenges

#### Sampling Methods and Metrics

The life detection papers catalogued in our survey represented a diverse array of life detection methodologies, geochemical parameters recorded, and extreme conditions which limit life (see Table 1 and Table 2). Differing methodologies which led to opposing life determinations for an environment required analysis to bridge the method gap in the opposing studies. To properly synthesize the information, one needs a thorough understanding of all methods used and the significance of data resulting from all experiments, an arduous task for a meta-analysis of meaningful sample size when each paper has different methods. Differing units between life detection methods were troublesome to compare in our statistical analysis and potentially introduced large errors. For example, salinity was reported with eight different units across the database, requiring density values for interconversion which were not always available. Further, authors often chose to record different geochemical factors depending on the site, their research background, funding, and the sampling year, which limits complete characterization and direct comparison of environment. Financial constraints, limitations of sampling technology, and access restrictions may influence life detection survey outcomes even before the researcher goes into the field.

**Table 2.** Life detection methodologies used by references included in the literature review have varying capabilities and limitations in targeting the known diversity of microbial taxa. Table design adapted from [52].

| Approach | Methods included in the literature review | All taxa targeted? | Non-viable or inactive cells included in analysis? |
| --- | --- | --- | --- |
| Culturing | Colony isolation, growth assay in extreme conditions, optical density of cultured samples, serial dilution plating | Yes, but selects against microbes that are yet unculturable | Yes |
| DNA and RNA | 16S rRNA sequencing, denaturing gradient gel electrophoresis (DGGE), DNA sequencing of genome or target gene, mRNA extraction | Depends on primers used. Archaea and/or eukaryotes are sometimes excluded from analyses | Yes, although obtaining sufficient biomass for a signal is difficult. mRNA can be used to isolate transcriptionally active cells and extracellular DNA is often an indicator of recent cellular activity. Extracellular nucleotides can be preserved in hypersaline brines<sup>53</sup> |
| Isotope Probing | [14C]-bicarbonate fixation, [13C]-enriched fatty acids, [3H]-leucine incorporation into proteins, [3H]-thymidine, [3H]-acetate incorporation, [15N] in NO<sub>2</sub>-, NO<sub>3</sub>-, and N<sub>2</sub>O, Dipicolinic acid triggered terbium ion [Tb3+] luminescence endospore viability assay, and other radioisotopic assays such as detection of methane production | Yes, although the choice of probe might exclude some organisms | Not typically |
| Metabolism | ATP detection, biochemical assays (e.g. oxidase, catalase, dehydrogenase), methane oxidation and sulfate reduction rates, protein electrophoresis, respiration assay of CO<sub>2</sub> | Yes, although the choice of assay might exclude some organisms | Depends. For example, ATP is widely considered to be an indicator of viable cells, but it is readily preserved and accumulated in hypersaline environments, and the enzyme-based luminescence assay is inhibited by ionic interference<sup>18, 20</sup> |
| Microscopy | Scanning electron, light | Yes | Yes |
| Molecular identification | Use of mass spectrometry, pyr-GC-MS, and GCMS to detect dissolved organic matter, PLFAs, proteins, polysaccharides, methane, ethane, and acetate | Usually, PLFA analysis excludes archaea, for example | Yes |
| Stain | Acridine Orange, BacLight, CARD FISH, DAPI, DTAF, DPA, FACS, Propidium Iodide, SYBR Gold, SYBR Green I with and without lysing | Usually, DPA excludes Archaea and Eukarya, for example | Yes, by BacLight, CARD FISH, DTAF, DPA, and FACS. Using SYBR Green I or Propidium Iodide without lysing cells will target vegetative cells, and with lysing cells, will target all cells. Otherwise, no |

| Dormant cells included in analysis? |  | References included in literature review |
| --- | --- | --- |
| With often-specialized resuscitation procedures, e.g. by heat shocking or freeze-thaw steps |  | 10; 12; 36; 41; 42; 43; 44; 46; 49; 50; 54; 55; 56; 57; 58; 59 |
| Depends on lysing procedure; Fast DNA Spin kit and mechanical disruption via bead beating lyses endospores as well as cells, whereas permeabilization and lysis achieved through lysozyme and achromopeptidase, for example, exclude spores. mRNA extraction does not include dormant cells in the analysis |  | 10; 29; 36; 42; 43; 44; 45; 46; 47; 48; 49; 50; 54; 55; 56; 57; 58; 59; 60; 61; 62 |
| No, although it can be if performed on cultured endospores after resuscitation. Dipicolinic acid triggered terbium ion [Tb3+] luminescence is the exception as an endospore viability assay |  | 29; 42; 48; 50; 56; 61; 62 |
| No |  | 25; 40; 41; 50; 55; 61 |
| Yes |  | 10; 25; 44; 62 |
| Yes, and mass spectrometry of whole cells and pyr-GCS-MS of dipicolinic acid, found only in endospores, can be used to isolate the dormant population <sup>63, 64</sup> |  | 25; 44; 45; 47; 49; 55; 57 |
| Yes, by BacLight, DTAF, DPA, and FACS. DNA stains such as SYBR Gold and SYBR Green I can detect spores if lysed with Fast DNA SPIN kit or mechanical disruption via bead beating. Acridine Orange and DAPI only stain endospores with specific treatment (heat, MAA, and ethanol) <sup>35</sup> |  | 10; 13; 14; 40; 41; 43; 44; 45; 47; 56; 60; 62 |

#### Cell Viability

There are myriad, highly diverse methods used to detect microbial life in extreme environments, but not all detection methods are equal in scope, precision, or intended target. Some life detection methods exclude unculturable microbes, archaea, or eukaryotes. Perhaps more importantly, some life detection methods differentiate between active microbial populations, dormant cells, and dead cells or fragments, while others do not.

Table 2 groups all life detection approaches used in publications for this database into seven categories. While culturing, isotope probing, metabolic assays, and some staining methods can separate viable cells from dormant, dead, or preserved remains, others do not. For example, microscopy without viability staining, the identification of molecular biomarkers, and nucleotide sequencing do not discern between viable and non-viable cells. Life detection in extreme environments using only these methods could result in false positives arising from preserved biological material.

#### Dormancy

The detection of dormant cells, particularly the specialized structures involved in dormancy such as endospores or cysts, is another challenge of life detection methodology. Dormancy is a reversible state of low metabolic activity in which cells can persist for extended periods without division^34^. Publications seldom incorporate dormant and slow-growing cells into the working definition of microbial life, thereby excluding these organisms from detection by using methods which detect active cell communities (Figure 5). Cultivation is still considered to be the gold standard for assessing the viability of spore-forming bacteria, but not all spores revive in laboratory conditions without specialized procedures such as heat shock or freeze-thaw steps. Unlike their bacterial counterparts, spores cannot be analyzed using fluorescent dyes such as acridine orange without specialized treatment^35^, though there are promising alternatives. Life detection studies that address dormancy can overemphasize endospores, thereby excluding from analysis the non-sporulating bacteria, archaea, and eukaryotes which are also capable of dormancy through the self-regulation of metabolism and growth rate. Excluding dormant organisms from life detection surveys may cause a false negative, in which both abundance and diversity reports of a microbial population are underrepresented.

**Figure 5.**
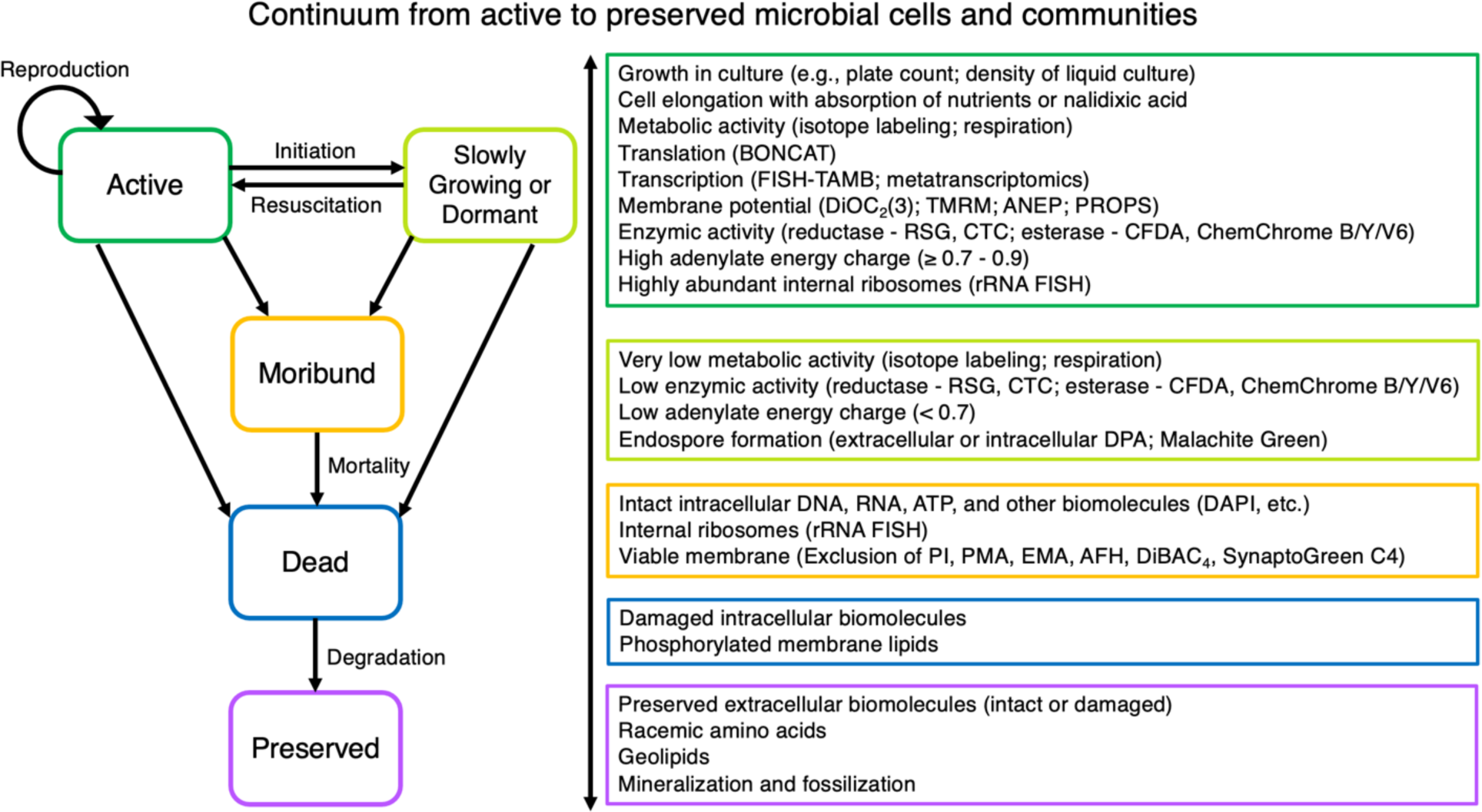
Microbiological methods target different populations along the continuum from active to preserved microbial cells and communities. At left, the physiological states within the continuum are defined by a level of activity or viability of a subpopulation of microbial cells. Methods shown at right (including specific examples) define the physiological state in the figure. For example, a moribund cell is one that has a viable membrane, and might have intact intracellular ribosomes or biomolecules. However, it lacks the traits of the active, dormant, or slow-growing cells above it; a moribund cell is unable to be revived and shows no evidence of cellular activity. Observations are placed at the lowest applicable activity but may occur, with confirmation for organisms at higher activity levels. Some placements are current topics of debate; phosphorylated membrane lipids are placed within the ‘Dead’ category, but could also be considered a part of ‘Moribund’. Microbial cells persist in the environment in all subpopulations defined within the continuum, but not every cell will become dormant, slowly growing, moribund, or preserved. Figure design adapted from [51] and content modified after T. Kieft & T.C. Onstott.

#### Experiments in situ vs. in vitro

In order to reproduce extreme conditions it is often necessary to oversimplify the complex conditions experienced by microbial communities in the field in order to test the limit of survival. This may misrepresent the true limit of survivability for organisms which have adapted to live under specific polyextreme conditions that may enhance or reduce habitability within a single extreme. Additionally, the absence of essential microbial community members or symbiotic partners and the timescale of the experiments may create false positives or negatives. Results for these experiments are greatly affected by the choice of model organism and the complexity of cultivation and experimental conditions in the lab. That said, laboratory experiments offer invaluable contributions for measurements that are difficult if not impossible to perform in certain field locations. Data analyses to follow (such as this study) need to take care in understanding though this is a proxy measurement, not an *in situ* measurement. A notable example from this study would be the limit for water activity from the Discovery Basin, Mediterranean Sea, that was revealed through subsequent lab experiments rather than in the environment.^36^

Beyond verifying findings from the field, researchers sometimes subject microorganisms to extreme conditions significantly past their environmental limits. For example, Kish et al.^12^ showed pressure tolerance in halophiles over six times higher than pressures found in any terran crustal environment, marking the “limit” of life as one that likely will never be reached in the field. While this can inform the potential habitability of other worlds, it is challenging to integrate with data from natural sites to understand polyextreme effects.

### Recommendations

#### Consider standardized methods and reporting protocols for natural samples at the limits of habitability

The difficulties our study encountered in comparing life detection surveys across global biomes could be alleviated by community conventions for methods and data reporting. While some discrepancy between studies is inevitable, greater standardization of life detection methodologies and sampling campaigns is feasible and highly valuable. Researchers working in extreme environments would be advised collect a standard suite of baseline physicochemical parameters during sampling. Only by standardizing methods can scientists publish results that are replicable and comparable for analog sites, enabling future investigators’ access to a broad range of data and increasing the credibility of published findings. In addition to scientific benefits, these standards could enhance return on investment for funding agencies and minimize repeat visits to field sites, thus promoting sustainability of rare or vulnerable environments.

Standard reporting protocols from journals and data portals would help ensure the comprehensive documentation of environmental physicochemical parameters and utilization of life detection methods that identify dormant and quiescent microbes, include phyla from all domains of life, and discern the viability of detected microbial populations.

#### Publish negative data and consider a centralized system

This study serves as a counterpoint to the positive publication bias rampant in the scientific literature. Positive results comprised 85.9% of all scientific literature published in 2007, but negative results are just as important to the scientific method^37^. The absence of negative results in the literature introduces bias into meta-analysis, subsequently misinforming researchers designing new studies. The underreporting of negative results also causes scientists to waste resources as they repeat experiments^38^. Negative life detections are often buried in the supplemental data sections of published life detection literature, or simply excluded from these papers. Bringing negative detections to the forefront will encourage other researchers to publish their negative results, as these are just as crucial to the advancement of science as positive findings.

This study would have been greatly improved by a standardized data collection and reporting system, such as those used for International Ocean Discovery Project cruises and other large-scale data collection projects. This system could be implemented across the field of geobiology to address data incongruity issues and reduce bias against the publication of negative data. Sites already exist for the publication of raw expedition data, such as PANGAEA, a member of the International Science Council World Data System. Useful tools such as built-in plotting capabilities already exist for this site, while sites in other disciplines, such as the Exoplanets Orbit Database at Exoplanets.org, allow for quick cross-publication analysis in the web browser to demonstrate general trends and easy educational demonstrations. Given the new emphasis on employing Open Science practices, the geobiology community is encouraged to make use of the centralized databases recommended for astrobiological research such as the NASA Astrobiology Habitability and Environment Database or the USGS Terrestrial Analog Data Portal. This system would increase useability of already-collected data, promote unit and measurement standardization (in conjunction with standardized methods), signal the value of negative results to the astrobiology community and encourage their documentation, and jump- start research into negative life detections by providing an easy-to-use dataset.

## Conclusion

This study proposed a new approach to understanding the limits of life. By cataloging negative life detections, this study sought to place an “outside limit” on the habitability ranges of life in polyextreme environments, complementing life detection findings that expand those limits outward (Figure 1). The lack of easily accessible and comparable data stymied attempts to use multivariate analysis to elucidate the driving polyextreme factors of negative life. While negative detections were found and recorded, the nature of the information that was published with these detections and the wide range of environments from which they were sourced complicated statistical analyses. Ultimately, understanding the confounding effects of polyextreme environments and experimental design decisions on life detection is just as important to enabling the search for life on other planets as determining a hypothetical habitable space. Improving life detection techniques to better quantify negative results can elucidate the limitations of commonly used biological assays and inform sampling procedures in both low- and high-biomass environments. Standardizing life detection methods and data reporting would improve data useability, save time and cost, and inspire more research into the existing troves of astrobiology data.

## METHODS

### Literature search

This study surveyed a wide variety of polyextreme habitats with negative life detection reports and documented all available data on factors that could influence habitability. These polyextreme environments have two or more exceptionally harsh properties and were targeted based on the assumption that the limits of life are most often encountered in these locations. The study utilized a network of related articles stemming from keywords specific to polyextreme biome genres.

To begin this study, a selection of extreme environment classes or “biome genres” were defined to establish a list of search terms^3^. These terms include “Hydrogeothermal,” “Polar Water,” “Desert Aridity,” “Deep Hypersaline Anoxic Basin,” and “Polar”, among others.

Descriptive words and the known locations of a given biome genre (Don Juan Pond in Antarctica, the Atacama Desert, etc.) were used as search terms on both Google Scholar and Web of Science. Additional keywords and referencing history were then drawn from articles that included negative life detections and used to extend the search. Papers were also sourced from discussions with colleagues and conference proceedings. An initial extensive article collection was performed in 2020 with a second article collection in 2023 based on the citation record of papers that recorded negative life detection events.

### Life Detection Terminology

For the purposes of this survey, the categorization of the presence or absence of life in each sample or sampling site is referred to as the ‘life status.’ A ‘life detection’ occurs when instrument data and available evidence support the conclusion that there is an active or dormant microbial community or even individual microbes in the sample. A reported absence of life in a sample or a negative result for an experiment looking for life is called a ‘negative life detection’ and may represent a vacant niche. Negative detections do not prove sterility of an environment, as the methods may not be sufficiently sensitive, broad, or qualified to detect organisms in that particular niche. The definition of a negative detection includes data within error of the noise floor of the detection method. Negative life detections can be used, however, to support hypotheses of sterility in combination with other analyses.

### Identifying negative results in literature review

Each article was examined for cases of reported or unreported negative life detections.

When a negative life detection was found, it was added to the site list and any available supporting data characterizing the site’s properties and the experimental methods were placed into a parameter database. This study analyzed over one hundred papers and found twenty publications with sites where at least one sample indicated a negative life detection (see SI for article list). In total, environmental datasets drawn from 28 publications were used for the construction of the parameter database. Negative results were discussed in some reports, but often did not include detailed negative data or metadata. In these cases, corresponding authors were contacted to retrieve data not explicitly included in the publication. Negatives not highlighted in a publication were more difficult to find and often required close reading of tables, charts, and supplementary data. For example, a difference in the number of field samples taken and analyzed samples noted in data tables often indicated a failed analysis or preparation, preferential analysis of a subset of samples, or omission of fully or partially processed negative results.

### Data standardization and normalization

In order to perform statistical analyses on the collected environmental parameters, the catalogued data was standardized with common units for measured properties. Data points were also removed from the database for parameters that did not meet the following thresholds: ≥ 5 non-repeated values recorded for the variable, values from ≥ 3 papers, and values from ≥ 10% of samples. After these exclusions, the environmental dataset consisted of 31 environmental variables and 398 observations from 28 individual papers (see SI).

The descriptive statistics such as the minimum, maximum, and standard deviation were gathered for each physicochemical parameter. These ranges were examined for anomalies and to understand the population of observations in the database. Statistical analyses (variance and standard deviation) revealed that the data for most variables was not normally distributed, representing the selection criteria for extreme environmental conditions. Imputation was attempted to address missing environmental observations, but due to the low percentage of samples with complete datasets in the database (48%) it was found to be ineffective. This eliminated methods such as Principal Component Analysis (PCA) from our potential set of analyses.

### Statistical methods

Non-metric multidimensional scaling (NMDS) and analysis of similarities (ANOSIM) were applied to understand the driving variables of the dataset. NMDS is a data dimensionality reduction method better suited to detecting clusters in data with many zeros as compared to PCA. NMDS requires a dataset of positive numbers and cannot include empty cells. To address the latter, all empty cells were filled with zeros. Then, for each column containing a negative number, the absolute value of the largest negative was added to all cells in the column (e.g., if the column contained {-5, -2, 4}, 5 would be added to each value, resulting in {0, 3, 9}). NMDS was completed on the full adjusted environmental dataset by the metaMDS function in the vegan R package using Bray-Curtis dissimilarity. The top three dimensions were extracted from the data, with 100 repetitions to find the best fit. The ANOSIM was carried out by the anosim function in the vegan package, and used Bray-Curtis dissimilarity was used to assess the extent to which the observations grouped by categories such as biome genre, location, and life status^39^.

Hypotheses regarding the effect of a single environmental variable on the occurrence of a negative detection were tested with binary logistic regression. A bivariate logistic regression was performed to ascertain the effects of certain parameters on the likelihood that life occurred in a sample. This analysis was performed using the Generalized Linear Model (GLM) function in the stats R package.

## ACKNOWLEDGEMENTS

The authors would like to thank Dr. Benedicte Menez for their insights which helped formulate this project. We also thank Jay Friedlander for his graphic design assistance in creating Figure 1. We acknowledge Michael Chesnes and Elissa Sperling for their assistance with the literature search, in particular Elissa’s assistance compiling literature in Russian. The authors gratefully acknowledge financial support from the Canadian Institute for Advanced Research Earth 4D: Subsurface Science and Exploration Catalyst grant program and the NASA Interdisciplinary Consortia for Astrobiology Research Award #80NSSC18K1140.

## AUTHOR CONTRIBUTIONS

Conceptualization: H.V.G. Framing and funding: H.V.G. and M.R.O. Methodology: H.V.G., L.E.R. and G.W.. Investigation: C.I.P., L.E.R., A.H.F. H.V.G. and F.N. Analyses: L.E.R. and A.H.F. with assistance from C.I.P., G.W. and F.N. Visualization: A.H.F., C.I.P. and L.E.R. Supervision: H.V.G. Writing—original draft: H.V.G., L.E.R., A.H.F. and C.I.P. Writing—review and editing: G.W., F.N. and M.R.O.

## DATA AVAILABILITY

Compiled data necessary to reproduce the analyses and figures for this paper will be made available with an archived DOI at the corresponding authors Zenodo.

