## Supplemental Information for "The Vacant Niche Revisited: Using Negative Results to Refine the Limits of Habitability"

**References included in the negative life detection database**

Papers with an asterisk were initially included but removed during the data cleaning process and thus not used in statistical analysis.


Belilla, J. et al. Hyperdiverse archaea near life limits at the polyextreme geothermal Dallol area. *Nat. Ecol. Evol.* **3,** 1552–1561 (2019).

Bryan, N. C., Christner, B. C., Guzik, T. G., Granger, D. J., and Stewart, M. F. Abundance and survival of microbial aerosols in the troposphere and stratosphere. *ISME J.* **13,** 2789–2799 (2019).

Cameron, R. E., Morelli, F. A., and Randall, L. P. Aerial, aquatic, and soil microbiology of Don Juan Pond, Antarctica. *Antarc. J. U.S.* **7,** 254–258 (1972).

Connon, S. A., Lester, E. D, Shafaat, H. S., Obenhuber, D. C., and Ponce, A. Bacterial diversity in hyperarid Atacama Desert soils. *J. Geophys. Res*. **112,** (2007).

Cragg, B. A., and Parkes, R. J. Bacterial profiles in hydrothermally active deep sediment layers from Middle Valley (NE Pacific), Sites 857 and 858. In Mottl, M. J., Davis, E. E., Fisher, A.T., and Slack, J.F. (Eds.) *Proc. Ocean Drill Prog., Sci. Results* **139,** 509–516 (1994).

Cragg, B. A., Summit, M., and Parkes, R. J. Bacterial profiles in a sulfide mound (Site 1035) and an area of active fluid venting (Site 1036) in hot hydrothermal sediments from Middle Valley (northwest Pacific). In Zierenberg, R. A., Fouquet, Y., Miller, D. J., and Normark, W.R. (Eds.), *Proc. Ocean Drill Prog., Sci. Results*, **169,** 1–18 (2000).

Davis, E. E. and Wang, K. Present and past temperatures of sediments at site 857, Middle Valley, Northern Juan de Fuca Ridge. In Mottl, M. J., Davis, E. E., Fisher, A.T., and Slack, J.F. (Eds.) *Proc. Ocean Drill Prog., Sci. Results* **139,** 565-570 (1994).

Drees, K. P. et al. Bacterial Community Structure in the Hyperarid Core of the Atacama Desert, Chile. *App. Env. Microbiol.* **72,** 7902–7908 (2006).

Fox-Powell, M. G., Hallsworth, J. E., Cousins, C. R., and Cockell, C. S. Ionic Strength Is a Barrier to the Habitability of Mars. *Astrobiology* **16,** 1–16 (2016).

Gemmell, R. T. Extremely halophilic archaea from ancient evaporite deposits. Doctoral dissertation, the University of Leicester (1996).

Goordial, J. et al. Nearing the cold-arid limits of microbial life in permafrost of an upper dry valley, Antarctica. *ISME J.* **10,** 1613–1624 (2016).

Hallsworth, J. E. et al. Limits of life in $MgCl_2$-containing environments: chaotropicity defines the window. *Env. Microbiol.* **9,** 801–813 (2007).

Heuer, V. B. et al. Temperature limits to deep subseafloor life in the Nankai Trough subduction zone. *Science* **370,** 1230–1234 (2020).

Inagaki, F., Nunoura, T., Nakagawa, S., Teske, A., Lever, M., Lauer, A. Biogeographical distribution and diversity of microbes in methane hydrate-bearing deep marine sediments on the Pacific Ocean Margin. *Proc. Natl. Acad. Sci. U.S.A.* **103,** 2815–2820 (2006).

*Kish, A., Griffin, P. L., Rogers, K. L., Fogel, M. L., Hemley, R. J., & Steele, A. High-pressure tolerance in halobacterium salinarum NRC-1 and other non-piezophilic prokaryotes. *Extremophiles* **16,** 355–361 (2012).

La Cono, V. et al. Unveiling microbial life in new deep-sea hypersaline Lake *Thetis*. Part I: Prokaryotes and environmental settings. *Soc. Appl. Microbiol.* **18,** 1–19 (2011).

Murray, A. E. et al. Microbial life at –13ºC in the brine of an ice-sealed Antarctic lake. *Proc. Natl. Acad. Sci. U.S.A*. **109,** 20626–20631 (2012).

*Navarro-González, R. et al. Mars-Like Soils in the Atacama Desert, Chile, and the Dry Limit of Microbial Life. *Science* **302,** 1018–1021 (2003).

*Peters, B., Casciotti, K. L., Samarkin, V. A., Madigan, M. T., Schutte, C. A., and Joye, S. B. Stable isotope analyses of $NO_2^-$, $NO_3^-$, and $N_2O$ in the hypersaline ponds and soils of the McMurdo Dry Valleys, Antarctica. *Geochim. Cosmochim. Acta.* **135,** 87–101 (2014).

Shipboard Scientific Party. Middle Valley: Bent Hill Area (Site 1035). In Fouquet, Y., Zierenberg, R. A., Miller, D. J., et al. *Proc. Ocean Drill Prog., Sci. Results*, **169,** 35-152 (1998).

Siegel, B.Z., McMurty, G., Siegel, S.M., Chen, J., and LaRock, P. Life in the calcium chloride environment of Don Juan Pond, Antarctica. *Nature* **280,** 828–829 (1979).

Steinle, L., Knittel, K., Felber, N., Casalino, C., de Lange, G., Tessarolo, C., et al. Life on the edge: active microbial communities in the Kryos $MgCl_2$-brine basin at very low water activity. *ISME J.* **12,** 1414–1426 (2018).

Takai, K., Moser, D. P., Onstott, T. C., Spoelstra, N., Pfiffner, S. M., Dohnalkova, A., et al. *Alkaliphilus transvaalensis* gen. Nov., sp. nov., an extremely alkaliphilic bacterium isolated from a deep South African gold mine. *Int. J. Syst.* **51,** 1245–1256 (2001).

Takai, K. et al. Cell proliferation at 122ºC and isotopically heavy $CH_4$ production by a hyperthermophilic methanogen under high-pressure cultivation. *Proc. Natl. Acad. Sci. U.S.A.* **105,** 10949–10954 (2008).

Tang, A. et al. 16S rRNA gene sequence analysis of halophilic and halotolerant bacteria isolated from a hypersaline pond in Sichuan, China. *Ann. Microbiol*. **61,** 375–381 (2011).

Tsiamis, G. et al. Prokaryotic community profiles at different operational stages of a Greek solar saltern. *Res. Microbiol.* **159,** 609–627 (2008).

Yakimov, M. M. et al. Microbial community of the deep-sea brine Lake Kryos seawater-brine interface is active below the chaotropicity limit of life as revealed by recovery of mRNA. *Env. Microbiol.* **17,** 364–382 (2015).

*Ziolkowski, L.A., Wierzchos, J., Davila, A.F., and Slater, G.F. Radiocarbon Evidence of Active Endolithic Microbial Communities in the Hyperarid Core of the Atacama Desert. *Astrobiology* **13,** 607–616 (2013).

**ANOSIM Results**

| **Grouping** | **ANOSIM Statistic R** | **Significance (p)** | **Number of Permutations** |
|---|---|---|---|
| Location | 0.9005 | 0.001 | 999 |
| Genre | 0.6381 | 0.001 | 999 |
| Reference | 0.893 | 0.001 | 999 |
| Life Status | 0.01344 | 0.132 | 999 |
